# Surviving Medical School During a Pandemic: Experiences of New York Medical Students During the Height of SARS-CoV-2

**DOI:** 10.1101/2024.06.18.599502

**Authors:** L. M. Knight, Divya Seth, David A. Zuckerman, Eli J. Rogers, Zain Talukdar, Robert Holloway, Caroline Gómez-Di Cesare, Mariluz Henshaw, Michael Privitera, Frank Dowling

**Affiliations:** University of Rochester School of Medicine, Rochester, NY, USA; Brown University, Providence, RI USA; Touro College of Osteopathic Medicine, Harlem, NY, USA; New York Medical College, Valhalla, NY, USA; Bassett Healthcare Network, Cooperstown, NY, USA; Renaissance School of Medicine, Stony Brook, NY, USA

**Author notes:** **Address correspondence to:** L. M. Knight, MD, Brown University, 593 Eddy Street, Aldrich Building, Room 126; Providence, RI, 02903. **Statements and Declarations:**. **Funding:** No funding was received for conducting this study. **Competing Interests:** The authors have no relevant financial or non-financial interests to disclose. **Author Contributions:** All authors contributed to the study conception and design. Material preparation, data collection and analysis were performed by L.M. Knight, Frank Dowling, Caroline Gómez-Di Cesare, Mariluz Henshaw, Divya Seth, and Eli Rogers. The first draft of the manuscript was written by Leanna Knight and all authors commented on previous versions of the manuscript. All authors read and approved the final manuscript.

**Keywords:** COVID-19, Medical Education, Medical Students, Medical Student Survey, MedicalSchools, Medical Licensing

## Abstract

**Background:** The COVID-19 pandemic dramatically altered the landscape of medical education. While patients overwhelmed hospital systems, lockdowns and social distancing recommendations took priority, and medical education was pushed online. Early in 2020, New York State (NYS) was hit especially hard by COVID-19.

**Objective:** This study sought to understand the effect of the COVID-19 pandemic on medical students well-being and education.

**Methods:** NYS medical students responded to a six-question survey during April and May 2020. Questions assessed self-reported changes in stress levels, academic performance, and board preparation efforts. Open-ended data was analyzed using a modified grounded theory approach.

**Results:** 488 responses across 11 medical schools were included (response rate of 5.8%). Major themes included: standardized test-related stressors (23%), study-related changes (19%), education and training concerns (17%), financial stressors (12%), and additional family obligations (12%).

Second year students reported more stress/anxiety than students in other years (95.9%, *p*-value< 0.00001). Reported stress/anxiety, effects on exam preparation, and anticipated academic effect varied by geographics.

**Conclusions:** While all NYS medical students reported being greatly affected, those closest to the NY City pandemic epi-center and closest to taking the Step 1 exam were the most distressed. Lack of flexibility of the medical education system during this public health emergency contributed to worsened student well-being. It is time to make plans for supporting the long-term mental health needs of these physicians-in-training and to examine ways the academic medical community can better adapt to the needs of students affected by a large public health emergency in the future.

## Introduction

Medical students have long been adversely affected by mental health challenges associated with medical education (1-6). These challenges for medical students were worsened by the COVID-19 pandemic; various short-term effects have been documented, including poor sleep quality and a high prevalence of depression, anxiety, and burnout (7). Currently, the long-term effect of the COVID-19 crisis on medical students, their careers, and their mental health remains unknown. Early in 2020, New York State (NYS) was hit especially hard by COVID-19 and the effects are still being felt today (8). Clerkship activities were paused, preclinical classes went online, and some medical students volunteered in the response effort (9). Early during the first wave of the COVID-19 pandemic, state medical societies received preliminary communications suggesting medical students may be experiencing unique stressors. Following up on these communications, the Medical Society of the State of New York (MSSNY) conducted a multi-institutional survey using closed and open-ended questions to understand the emotional and educational effect of the COVID-19 pandemic on NYS medical students.

In this study, we aimed to assess the effects of the COVID-19 pandemic on medical students in NYS, one of the first epicenters of the COVID-19 pandemic in the United States. We took this opportunity to provide an analysis of medical student education and wellness data from the MSSNY survey conducted in NYS at the height of the first COVID-19 wave and consider the information gleaned from this survey to advocate for learner needs. Given our findings, we suggest that there are additional areas of concern to be addressed (1).

## Methods

The six-question survey was distributed to all 11 active MSSNY medical school chapters between 04/14/2020-05/02/2020. Medical student respondents of all years attending an allopathic or osteopathic medical school in NY were included in the study.

The survey was composed of closed and open-ended questions including:

1. During the COVID-19 pandemic are you feeling more stress/anxiety?
2. Do you feel you will be able to maintain or improve your performance from last semester?
3. Do you feel that this pandemic has interfered with your ability to prepare for Step or Shelf exams?
4. If you answered the previous questions, please elaborate on how this has impacted you.
5. If given the time and freedom from your current medical school responsibility, would you be active in the response efforts to COVID-19?
6. Would you like to share anything else?

Data was accessed by the team in early 2021. Each of the open-ended responses were analyzed by 2 assigned independent researchers (Knight, Dowling, Gómez-Di Cesare, Henshaw, Seth, Talukdar, or Rogers) using modified grounded theory with an inductive approach implementing the techniques described by Strauss and Corbin (10-12). Our analytic process featured six steps: 1. data “fractured” or “broken down into granular codes” during independent review; 2. incident-to-incident coding; 3. peer debriefing at each step; 4. focused coding of fractured codes into coherent subsuming categories or “themes” by two independent researchers; 5. axial coding through identifying relationships between codes and categories; and 6. production of findings (13). In the final analysis, only themes that were identified by both independent researchers were included. To minimize bias during the coding process, we blinded the name and location of each medical school. Researchers used the documented reflective process necessary for a rigorous application of grounded theory to open-ended responses.

## Results

488 student responses across 11 medical schools were included, with a response rate of 5.8%. Table I shows the major themes identified in open-ended responses. These include the effect of standardized test-related stressors (23% of respondents, n=113), study-related changes (19%, n=93), concerns about the quality of education and training (17%, n=82), financial stressors (12%, n=59), and additional family obligations (12%, n=52). Table II summarizes the responses to the six survey questions. Results were analyzed by school year with students in all years reporting increased levels of stress/anxiety. Second year students (class of 2022) reported near-universal increases in stress/anxiety. More second year medical students reported that the pandemic had interfered with their ability to prepare for Step or Shelf exams than medical students from other years. Geographic differences were seen across responses. Students from downstate NY, including NYC and the surrounding area, reported higher rates of stress/anxiety, more difficulties in maintaining or improving academic performance, and more difficulties preparing for standardized exams.

**Table 1.** Differences Based on Medical School Year or School Location.

| Table I. Differences Based on Medical School Year or School Location |  |  |  |  |  |
| --- | --- | --- | --- | --- | --- |
|  | Class of 21 | Class of 22 | Class of 23 | Class of 24 | P-Value |
| Difficulties Preparing for Step/Shelf Exams | 18.8% | 93.3% | 70.5% | 6.7% | <0.01 |
| Believe I cannot maintain/improve academic performance | 42.3% | 62.2% | 57.7% | 43.7% | <0.002 |
| Experiencing stress/anxiety | 95.9% | 73.8% | 82.6% | 66.7% | <0.00001 |
|  | Downstate (NYC and surrounding areas) | Upstate (the rest of NY State) |  |  | P-Value |
| Difficulties Preparing for Step/Shelf Exams | 65.4% | 61.4% |  |  | 0.000269 |
| Believe I cannot maintain/improve academic performance | 38.1% | 52.6% |  |  | 0.001258 |
| Experiencing stress/anxiety | 88.8% | 81.1% |  |  | 0.016629 |

**Table 2.** Medical Students Concerns During the Early Stages of the COVID-19 Pandemic.

| Table II. Medical Students Concerns During the Early Stages of the COVID-19 Pandemic |  |  |
| --- | --- | --- |
| Theme | Percent Reporting | Example |
| Step and Shelf | 28% (137) | Prometric has canceled my appointment for taking Step so I have no clue when I'll be allowed to take the exam. Because of this, I don't know how to formulate a study plan or if I should even be studying for Step at all. Although I'm still trying to study, my anxiety is making it incredibly difficult to focus on any of the material. |
| Studying Related Changes | 20% (97) | I am home, which is nice for the most part. But my family of 4 lives in a 1 bedroom apartment so finding time to study has been a horrible struggle. I've read articles about how quarantine is exposing students to their inequitable experiences. And that is my life right now. Unlike my friends, I don't have any personal space or any privacy to study effectively. I don't have the gift of silence or a set table and chair to my name to help me focus. I am behind and it is difficult to disclose to anyone, it's embarrassing. |
| Stress or Anxiety | 18% (89) | The uncertainty of everything has created massive anxiety |
| Quality of Education or Uncertainty about Future Training or School Communication | 20% (100) | As a third year medical student, concerns of insufficient training are at the forefront of my concerns. ... As it stands it seems we lost significant core clinical experience in our third year which will hopefully be made up in fourth year, but should the situation escalate again, likely will not be compensated for in our education. |
| Financial Negative Effect Reported | 14% (67) | My parents help pay for my school and apartment. My dad is currently unemployed as a result of the pandemic. |
| Family Obligations: Other | 10% (50) | Multiple sick friends/family that needed assistance, increased at home demands/duties |
| COVID-19 Fear | 8% (37) | I'm worried that I'm a carrier of the virus and will get my parents sick. A lot of people their age die and it makes me anxious. |
| COVID-19 Illness | 4% (12) | I've had multiple deaths in my immediate family from COVID19, financial stress for funerals, and family members sick with Covid19 hoping to recover. I have high anxiety of being infected myself. It's an emotional, mental, and physical strain on me. All I have in my mind is the well-being of my family. |
| Volunteering Desire Distress | 7% (33) | it's a strange kind of uselessness, being a medical student in a pandemic, particularly during the dedicated period of study for Step 1 - I feel that any time not spent studying is wasted but I also really want to volunteer and help more. |
| Family Obligations to Children | 2% (10) | Also had my time taken up by tutoring family members that are no longer in school and doing virtual learning, which has not seemed to be doing much for many NY school attending students. |
| Isolation or Loneliness | 2% (10) | My mental health has been struggling from the self isolation |
| Positive Academic Experience | 2% (10) | The pandemic has paradoxically given me more time to study during this crucial semester before Step 1/Level 1. Without the commute and extra distractions of leaving the apartment, I have had more time to thoroughly review my school course materials as well as board exam-specific tools. With only my spouse and I at home, I did not acquire extra responsibilities. |
| Acknowledges Privilege | 2% (8) | I'm fortunate to have few obligations and that my family is in good health. |

## Discussion

No one in medical education could have predicted at the beginning of the 2019-2020 academic year that the year would end with a country-wide shut down, over one million deaths, and our medical institutions being stressed beyond anything ever seen before (14). This data demonstrates that those who were in medical school during the pandemic faced many acute stressors within the context of social disruption and isolation. A sizable subset of students faced additional family obligations, financial pressures, and isolation. The uncertainty and late cancellations of Step and COMLEX licensing exams were major stressor and led many students to experience multiple or extended dedicated study periods. Delays inevitably required not only more time away from their medical education, but exacerbated severe financial stressors and social isolation. For these reasons among others, it is paramount that we take advantage of lessons learned.

As the pandemic has evolved, medical education has adapted. The once-rigid expectations on which the foundation of medical education is built have become more flexible and accommodating. Students can attend online lectures and more easily take sick days. Schools have become more proactive with COVID-related communications. The NBME and NBOME eventually developed ways to expand their exam testing capacity by relaxing requirements to allow medical schools to administer the Step exams.

In April of 2022, ACGME, AAMC, AACOM and ECFMG jointly published “Transition in a time of disruption. Practical Guidance to Support Learners in the Transition to Graduate Medical Education” as an anticipatory guide for residency programs and students as some of the hardest-hit students prepared to enter residency in the summer of 2022. The toolkit lays an important foundation for residents who had their medical education disrupted during medical school by the pandemic. While competency, training, and assessment tools likely will help to bridge the gap to assist pandemic-affected students to become competent residents, we are concerned about the long-term consequences of the unique and extraordinary stress medical students in the epicenter of the pandemic faced as this may contribute to increased vulnerability to the impact of severe stressors in the future. Going forward, increases in institutional flexibility and education disruption planning will be needed to protect and assist the next generation of physicians. Since physicians are needed on the front lines of any public health emergency, contingency planning to address disruptions in education of affected students is paramount.

## Conclusion

In this study, we demonstrated that the COVID pandemic was a great social disrupter that produced noteworthy negative effects on the medical students of NYS during the first wave, causing psychological distress and creating gaps in education that may leave many of these students behind their peers as they enter residency. It is imperative that medical educators not only develop plans to assist those most adversely affected by the pandemic so they can address potential training gaps to allow for a smoother transition into residency, but also develop disaster preparedness plans that include strategies to identify, address and mitigate the potential adverse mental, educational, social and financial ramifications on medical students and training physicians.

## Acknowledgments

We would like to thank MSSNY and its staff for their role in making this research more robust.

